# HOMINID: A framework for identifying associations between host genetic variation and microbiome composition

**DOI:** 10.1101/081323

**Authors:** Joshua Lynch, Karen Tang, Sambhawa Priya, Joanna Sands, Margaret Sands, Evan Tang, Sayan Mukherjee, Dan Knights, Ran Blekhman

## Abstract

Recent studies have uncovered a strong effect of host genetic variation on the composition of host-associated microbiota. Here, we present HOMINID, a computational approach based on Lasso linear regression, that given host genetic variation and microbiome composition data, identifies host SNPs that are correlated with microbial taxa abundances. Using simulated data we show that HOMINID has accuracy in identifying associated SNPs, and performs better compared to existing methods. We also show that HOMINID can accurately identify the microbial taxa that are correlated with associated SNPs. Lastly, by using HOMINID on real data of human genetic variation and microbiome composition, we identified 13 human SNPs in which genetic variation is correlated with microbiome taxonomic composition across body sites. In conclusion, HOMINID is a powerful method to detect host genetic variants linked to microbiome composition, and can facilitate discovery of mechanisms controlling host-microbiome interactions.

**Availability and implementation:** Software, code, tutorial, installation and setup details, and synthetic data are available in the project homepage: https://github.com/blekhmanlab/hominid.

Real dataset used here is from Blekhman et al. (Blekhman et al. 2015); 16S rRNA gene sequence data and OTU tables are available on the HMP DACC website (www.hmpdacc.org), and host genetic data are deposited in dbGaP under project number phs000228.

## Background

The microbial communities found in and on the human body are influenced by multiple factors (Consortium, Human Microbiome Project 2012). In addition to the clear effect of environmental factors on the microbiome, there is growing support for an impact of host genetics (Goodrich, Davenport, Waters, et al. 2016; Morton et al. 2015). Several candidate gene studies have found correlation between human genetic variation and the structure of the microbiome (Tong et al. 2014; Khachatryan et al. 2008; Knights et al. 2014). In addition, genome-wide approaches can also be useful to identify human genetic impact on the microbiome (Goodrich et al. 2014; Blekhman et al. 2015; Goodrich, Davenport, Beaumont, et al. 2016; Davenport et al. 2015). For example, Goodrich et al. used hundreds of twin pairs to calculate the heritability of the gut microbiome, and identify bacterial taxa that are heritable, such as Christensenellaceae (Goodrich et al. 2014). Researchers have also utilized quantitative trait locus (QTL)-mapping approaches in the laboratory mouse and have identified multiple loci associated with the structure of gut microbial communities, some of which overlap genes involved in immune response (Benson et al. 2010; Leamy et al. 2014). Moreover, studies have used joint human genetic variation and microbiome data to find associations between loci in the human genome and microbial taxa (Blekhman et al. 2015; Davenport et al. 2015; Bonder et al. 2016; Turpin et al. 2016). In our recent study, in addition to showing that human genetic variation is associated with the structure of microbial communities across ten body sites, we have identified human single nucleotide polymorphisms (SNPs) associated with variation in the microbiome, and found that these loci are highly enriched in immunity genes and pathways (Blekhman et al. 2015). This approach, which includes the joint analysis of host genetic variation (SNPs) and microbiome taxonomic composition data (usually an OTU table), has the important advantage of identifying specific host genes and pathways that may control the microbiome, thus shedding light on the biological mechanisms of host-microbiome interaction, and pinpointing potential disease-causing pathways. However, this analysis is complicated by the fact that the microbiome contains many taxa that can be used as potential molecular complex traits in the GWAS analysis. Testing many taxa reduces the power and multiple hypothesis testing correction makes the identification of associations challenging.

Here, we propose a framework for identifying host SNPs associated with microbiome composition using Lasso regression, named **HOMINID** (**Ho**st-**M**icrobiome **In**teraction **Id**entification; see **Figure 1** and **Supplementary Information**). Our method has several advantages: (1) it takes as input host genetic variation data (in a modified VCF format) and microbiome composition data (as an OTU table), to facilitate a simple analysis pipeline with no need to make new data formats; (2) HOMINID uses Lasso regression, which is specifically designed for cases where a relatively small number of taxa are correlated with host SNP genotype, as opposed to existing methods that use all taxa abundances; and (3) HOMINID uses stability selection with randomized Lasso to identify the specific microbial taxa that are correlated with each associated SNP.

**Figure 1.**
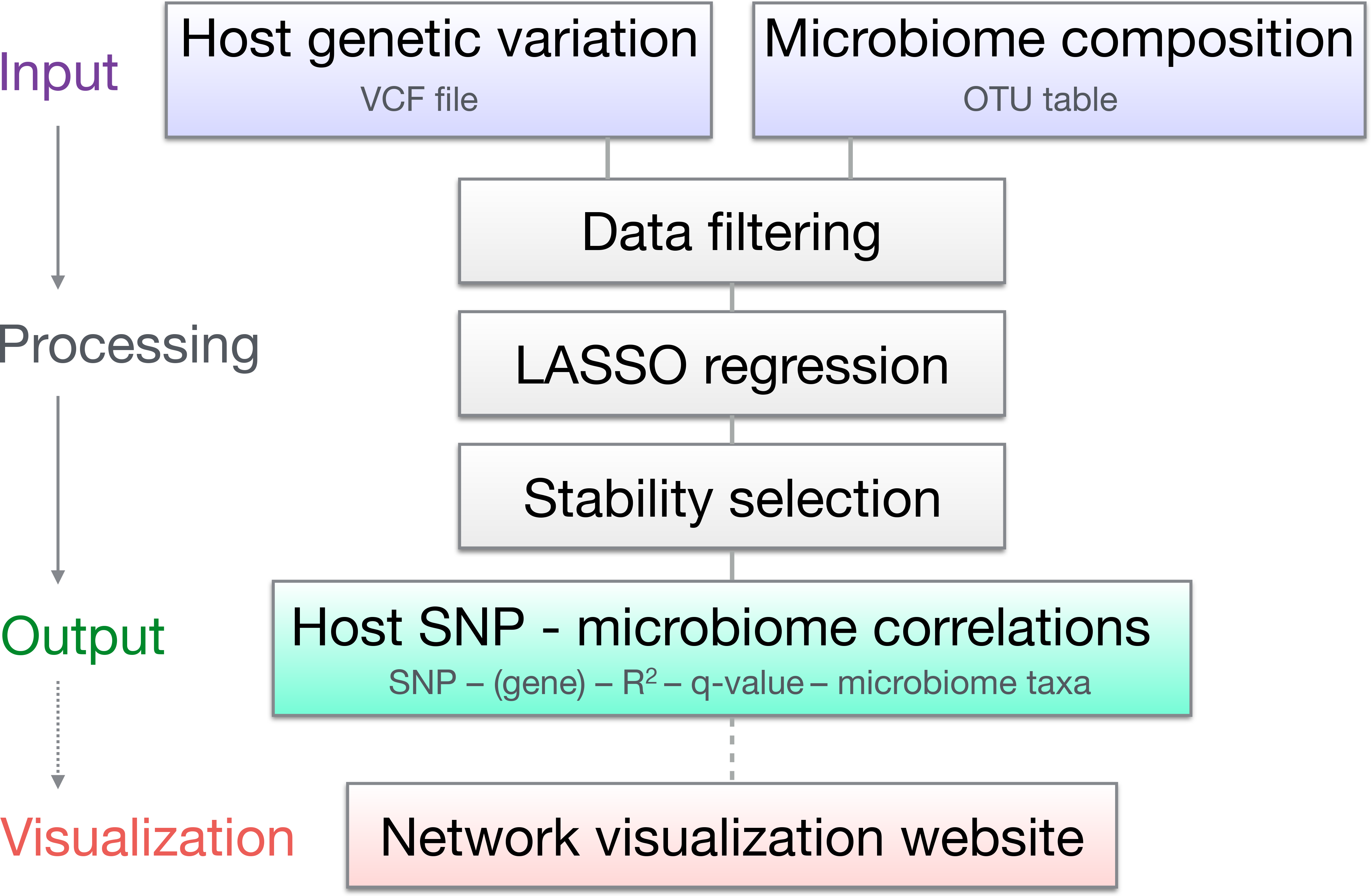
Illustration of the HOMINID pipeline.

## Materials and Methods

### HOMINID implementation

We implemented Lasso regression with the taxon relative abundances (arcsin sqrt transformed) as predictors and genetic variation at each SNP as response, for the purpose of identifying an additive effect between host genotype and microbiome features (see **Supplementary Information** and **Figures S1-S3**). In most situations, we expect at most a few taxa’s abundances to correlate with a SNP, therefore ordinary least-squares (OLS) regression, which includes all taxa abundances as predictor variables, might not be an appropriate model. Instead, we need a regression algorithm that selects only the few predictors (taxa) that correlate to host genetics and discards the rest. The Lasso linear regression model used for HOMINID is similar to OLS regression, except that it includes an additional penalty term that shrinks most regression coefficients to zero, resulting in a sparse solution; thus it predicts only a few taxa to correlate with the host genetics. The Lasso regression was implemented using the Python (version 2.7/3.5+) machine-learning library scikit-learn (Pedregosa et al. 2011), with microbiome relative abundances as predictors and SNP genotype as response variable. The penalty term was tuned via a five-fold cross-validation. How well the host genetics correlates with the microbiome is measured with the coefficient of determination, 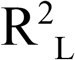. 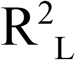 is the median R^2^ from five-fold cross-validation, with 100-times resampling. Also outputted are 95th percentile bootstrap confidence intervals from 10,000 bootstrap samples. Detailed description of the implementation of Lasso regression is available in the Supplementary Information.

### Identifying correlated SNPs and taxa

To identify SNPs that are predicted correlated to the microbiome (prediction positive) from the uncorrelated (prediction negative) HOMINID uses a q-value cutoff, which puts an upper bound on the False Discovery Rate (FDR). A cutoff value, 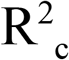, of 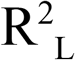is chosen such that the q-value, q(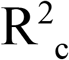), is equal to 0.1. A given SNP is predicted positive (predicted correlated to the microbiome) if 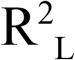≥ 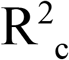. q(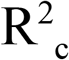) is determined by a permutation test, whereby for each SNP the sample labels are shuffled and Lasso regression is rerun ten times. q(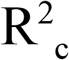) is defined as the fraction of permuted SNPs predicted positive divided by the fraction of unpermuted SNPs predicted positive (Subramanian et al. 2005). 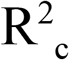is chosen such that q(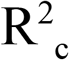) = 0.1. The taxa that are most strongly associated with a SNP are identified using Stability Selection with randomized Lasso (Meinshausen and Bühlmann 2010). Briefly, stability selection perturbs the regression coefficients and the penalty term in the Lasso regression, and then reruns the regression thousands of times. If the same predictors (taxa) are repeatedly selected, even when the odds are against them, then they are robust predictors. Full details on this procedure are available in the Supplementary Information.

### Controlling for other (non-taxon) covariates

HOMINID allows for controlling for any additional covariates (other than the microbiome) by including the covariates in the microbiome taxonomic table. This enables controlling for potentially confounding factors, such as individual age and sex. It also enables controlling for ancestry (or population substructure) by including the principal components (PCs) of the genetic variation data (Price et al. 2006; Pritchard et al. 2000). in the analysis. We performed two analyses, one including host genetic PCs as covariates (results in Supplementary Table S1), and one without these covariates (Supplementary Table S2). We excluded from the results SNPs for which there is a strong correlation with sex.

### Synthetic datasets

To test the performance of HOMINID we generated several synthetic datasets. “Taxon” absolute abundances (“counts”) were drawn from a log-series distribution. The log-series distribution is frequently used to represent species abundances (see, e.g., (Baldridge et al. 2016)), and it allows a range of abundances that spans several orders of magnitude, mimicking both rare and abundant taxa. Often in real abundance tables a large fraction of taxa have an abundance of zero (taxon either not present or not detected). The log-series abundance tables also had this quality; in our synthetic data, 21% of abundances are count zero. Synthetic SNP data were generated such that, for each SNP, N_ctc_ random taxa’s abundances correlate with that SNP’s genotype. Uncorrelated SNPs were created by permuting the sample IDs, preserving the minor allele frequency. Effect size was varied by adding “noise” to the SNP genotype data. Once the SNP and taxon-abundance data were generated, a measure of the effect size was calculated: the coefficient of determination, 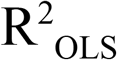, for an ordinary least square (OLS) multiple regression between the correlated taxa’s abundances and the SNP genotype. Since 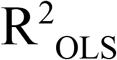is a characteristic of the input data before analysis by HOMINID, we call it the “input R^2^” to distinguish it from the R^2^ output by the HOMINID Lasso regression (aka the “output R^2^” or 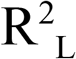). To examine data sets with smaller effect sizes, “noise” was added to the SNP data by swapping the genotypes of pairs of samples, reducing the correlation between the N_ctc_ correlated taxa and the host SNP genotype. Several data sets were created with progressively more “noise”, until 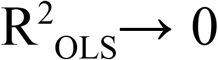. We created three sets of synthetic data to examine the performance of HOMINID on different qualities of the input data: Data set MAF varies the minor allele frequency, with MAF ranging from 0.10 to 0.50; data set CTC varies the number of correlated taxa from five to twenty; and data set TC varies the total number of taxa in the taxon table from 100 to 500. All data sets contain 500 SNPs each. Data in sets MAF and CTC comprise 1000 individuals; data sets in set TC contain 100 individuals. Data sets MAF and TC all have three correlated taxa per SNP. The MAF for data sets CTC and TC is 0.30.

### Human Microbiome Project data

In addition to the synthetic datasets described above, we also tested our method on a real dataset that includes both human genetic and microbiome data (Blekhman et al. 2015). This dataset includes 93 individuals for whom microbiome was profiled as part of the Human Microbiome Project, and for which host genetic variation information was extracted from shotgun metagenomics sequence data as described previously (Blekhman et al. 2015). We annotated the previously described set of 4.2 million high-quality single nucleotide polymorphisms (SNPs) using ANNOVAR (Wang, Li, and Hakonarson 2010) and focused the analysis on a set of 32,696 protein-coding SNPs. We further filtered this set to include only SNPs with minor allele frequency of at least 20% and SNPs for which we had data for at least 50 individuals. The number of SNPs actually tested varies across body sites, ranging from 12,400 to 14,651 SNPs, with a mean of 14,023. For the Stool microbiome data, which included 107 total taxa, running HOMINID on 14,469 SNPs using 12-core Intel Xeon E5-2680 2.50 GHz processors took 16 cpu hours.

### Comparison to other methods

The PERMANOVA (Anderson 2001; McArdle and Anderson 2001) analysis was done in R with the adonis function in the vegan (Oksanen et al. 2007) package. The model formula has the SNP genotype as numeric (not factor) predictor variables and the arcsin-sqrt transformed taxon relative abundance table as response variable. The method used to calculate pairwise “distances” was the default Bray-Curtis. The MiRKAT (Zhao et al. 2015) analysis was performed using the MiRKAT package in R. The Bray-Curtis dissimilarity matrix was computed on the arcsin-sqrt transformed taxon table. The matrix was then converted to a kernel matrix, and MiRKAT invoked for each SNP. Since both PERMANOVA and MiRKAT output p-values as measures of how well the taxon abundances correlate with each SNP’s genotype (whereas HOMINID outputs 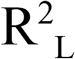values) we chose a cutoff value of p-value such that q(p_c_) = 0.1 to separate the prediction positives (correlated) from the prediction negatives (uncorrelated), much in the same way we chose the cutoff 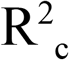to separate prediction positive/negative such that q(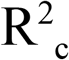) = 0.1 for the Lasso regression.

## Results

### Analysis using synthetic data

To assess HOMINID’s performance, we first used the pipeline on a comprehensive set of synthetic datasets (described above and in the Supplementary Information). These datasets were designed to simulate variation in several important factors, such as variation of the strength of correlation (the input R^2^) of the associated SNP with microbiome composition, variation in minor allele frequency (MAF) of the associated SNP, noise level in microbiome data, and the number of taxa associated with the SNP. After analyzing each of the datasets we calculated and plotted the method’s sensitivity, specificity, precision, negative predictive value (NPV), false positive rate (FPR), false negative rate (FNR), false discovery rate (FDR), and accuracy, as a function of the input R^2^, highlighting the effects of the variable factors above (see **Figs. 2A–D**, Supplementary Information and Supplementary Figures S4 - S43).

We found that the strength of correlation (input R^2^) between SNP genotype and the correlated taxa has little effect on HOMINID’s ability to identify the SNP, unless the correlation is very low (**Figs. 2A and 2B**, Supplementary Information, and Supplementary Figures S4 - S11). HOMINID achieved high sensitivity and specificity for R^2^ values of above ~ 0.05. The False Discovery Rate (FDR) is below 0.1 by design, and variation in FDR is due to imprecision (finite number of significant digits) in calculation of 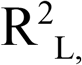and therefore imprecision in calculation of q. (**Figs. 2C** and **2D**). Similarly, variation in MAF does not affect HOMINID’s sensitivity, as data sets with different MAF follow the same behavior (**Fig. 2B**).

**Figure 2.**
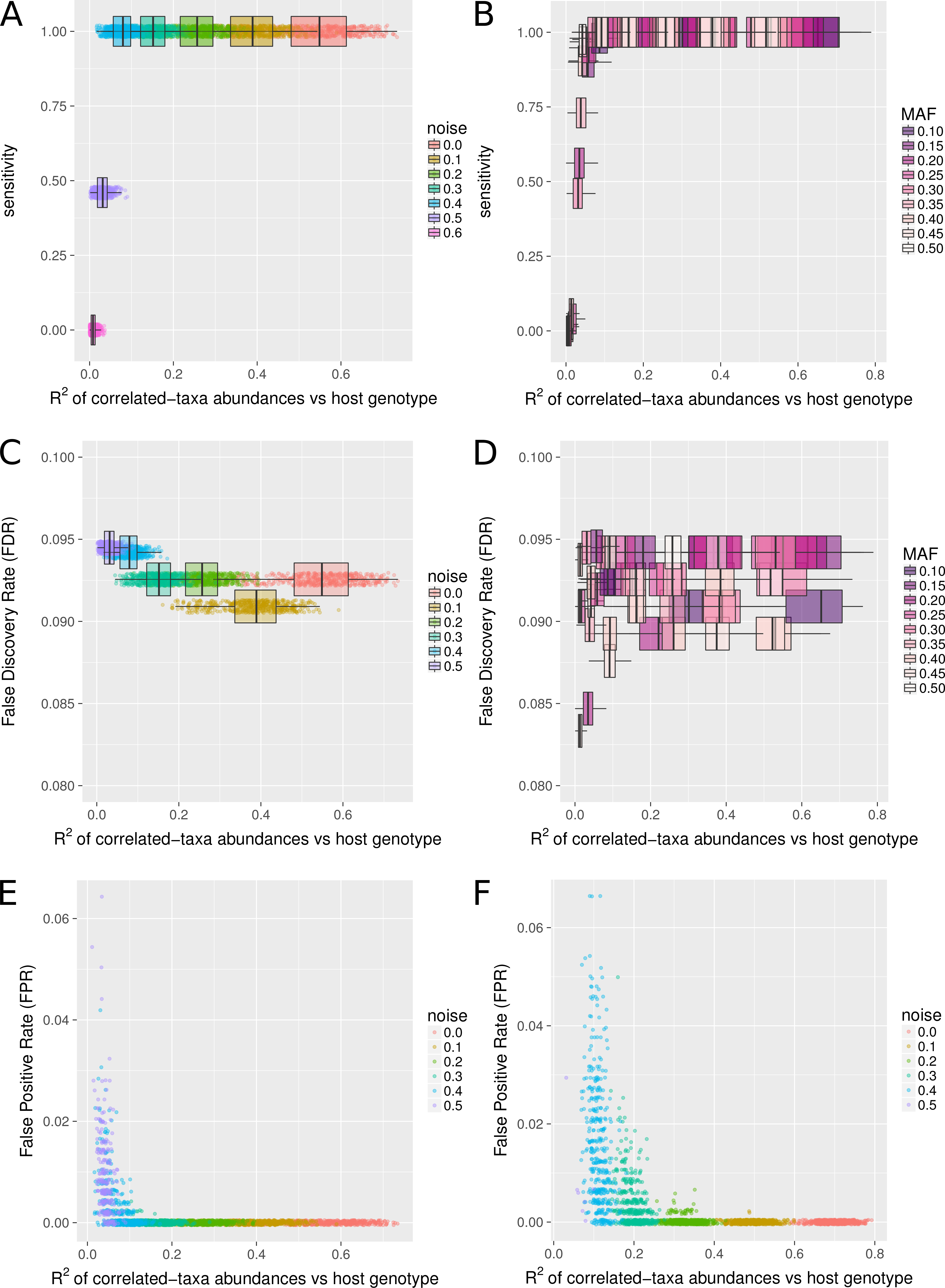
Assessment of HOMINID’s performance using synthetic data. Panels **A-D** assess how well HOMINID predicts the SNPs whose genotypes correlate with microbiome abundances, and panels **E** and **F** assess how well HOMINID predicts the specific taxa correlated with an associated SNP. **(A)** Sensitivity as a function of effect size (input R^2^) for the data sets with MAF=0.30. Different colored points and boxplots represent data sets with different noise levels and therefore different effect sizes. (**B**) Same as A with variation in input data MAF values represented by different colored boxplots. (**C)** FDR as a function of effect size (input R^2^) for data sets with just MAF=0.30. (**D**) Same as C with variation in input MAF values represented by different colored boxplots. (**E)** FPR for the stability selection step (identifying the taxa that associate with a SNP’s genotype), as a function of effect size (input R^2^) for data sets with three correlated taxa. (**F**) Same as E but with twenty correlated taxa.

One of HOMINID’s unique features is the ability to identify the taxa that are correlated with an associated SNP. We found that this prediction performs well, with accuracy approaching 1 and a false positive rate (FPR) of 0 for input R^2^ values larger than about 0.1, but drops off at lower R^2^ values (**Fig. 2E** and **Fig. 2F**, Supplementary Figures S26 and S27). The number of correlated taxa had a noticeable effect, whereby SNPs that correlated with more taxa had higher FPR (compare **Fig. 2E** with **Fig 2F**), although in all test datasets’ FPR remained < 0.07.

### Comparison to other methods

In order to assess HOMINID’s performance, we compared it to PERMANOVA (Anderson 2001; McArdle and Anderson 2001) and MiRKAT (Zhao et al. 2015), two platforms that can be used to identify host SNPs associated with microbiome composition. We note that HOMINID has a unique feature allowing it to identify the specific microbial taxa associated with each SNP. Since other approaches lack this option, the comparison centered around the ability to detect SNPs that are correlated with the microbiome, and not on the detection of correlated taxa. Our analysis included input datasets with various input R^2^ values and noise levels (various effect sizes), and compared the sensitivity of each method to detect the associated SNPs. We found that for median input R^2^ values (correlation between associated SNP and microbiome composition) of about 0.15 or above the three methods are all highly sensitive (**Fig. 3**). However, for lower input R^2^ values, HOMINID is more sensitive. Specifically, for the data set with median input R^2^ = 0.08 HOMINID’s sensitivity is 1, while the sensitivity of MiRKAT and PERMANOVA is 0.19 and 0.29, respectively (**Fig. 3**). Similarly, for median input R^2^ = 0.03 HOMINID’s sensitivity is 0.46, while the other methods’ sensitivities are 0.

**Figure 3.**
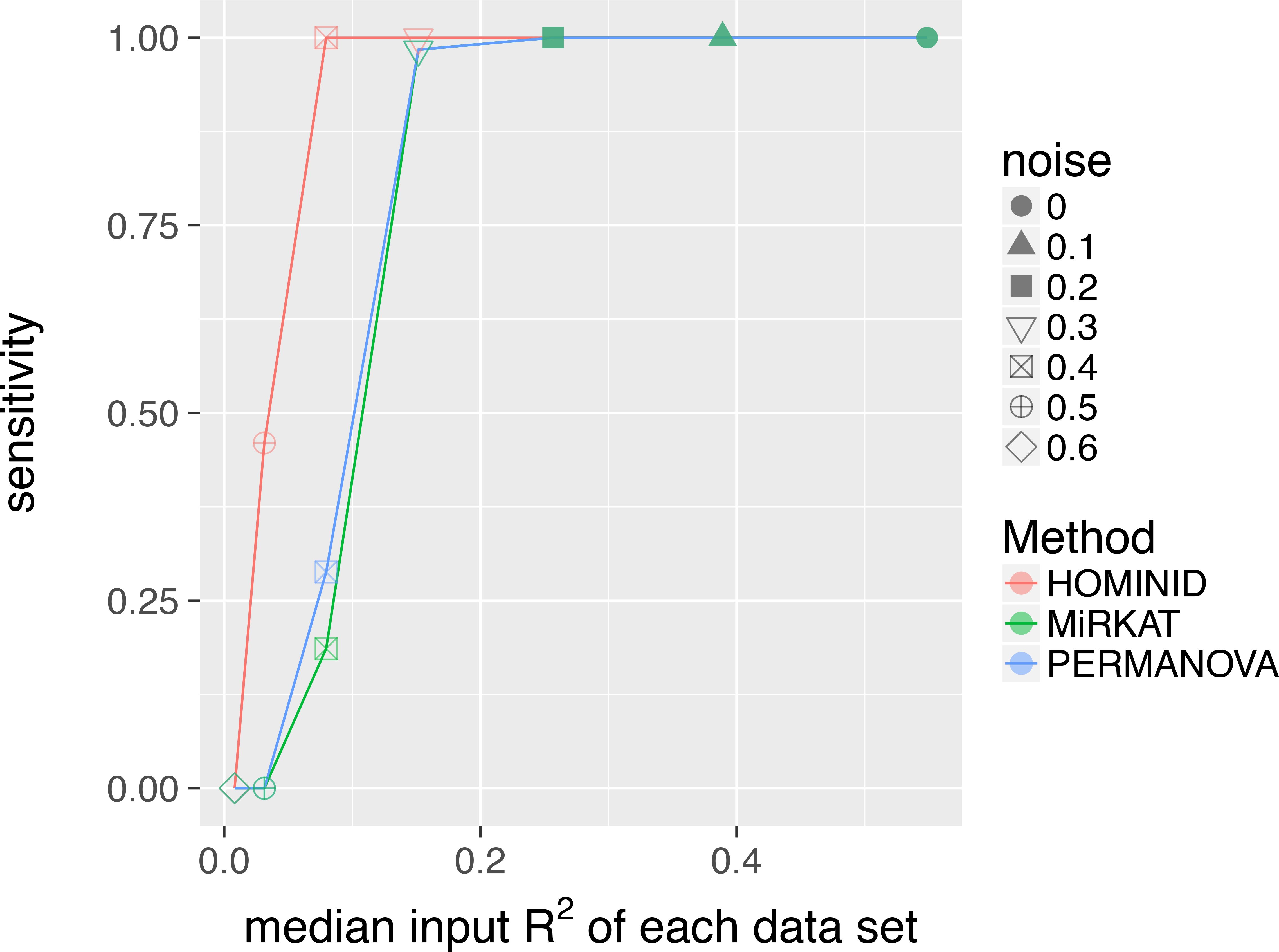
Comparison of the performance of HOMINID versus MiRKAT and PERMANOVA. Sensitivity is plotted as a function of effect size (input R^2^) for HOMINID (red), MirKAT (green), and PERMANOVA (blue). At high input R^2^ all three methods perform well, finding all SNPs that correlate with the microbiome. However, at smaller effect sizes (lower input R^2^), HOMINID is more sensitive.

### Human Microbiome Project data

We ran the HOMINID pipeline on a previously published data of microbiome and host genetic variation from the Human Microbiome Project cohort (Blekhman et al. 2015). We focused our analysis on coding SNPs with minor allele frequency ≥ 0.2, and identified SNPs for which permutation-based q-value ≤ 0.1 and the 95th percentile confidence interval for R^2^ does not include zero. To account for population substructure, we ran a second analysis including the genetic principal components (PCs) as additional covariates (Price et al. 2006; Pritchard et al. 2000). This resulted in the identification of 11 (regression with genetic PCs as covariates) and 6 (regression without genetic PCs) for a total of 13 unique associations between host SNP and microbiome composition across 15 body sites (see Supplementary Tables S1 and S2, respectively). As can be seen in Figure 4, HOMINID is able to detect SNPs with the expected pattern of association between host genetic variation and the microbiome. For example, for SNP rs2297345 in the gene *PAK7* we detected a correlation between genotype and a single microbial taxon, Propionibacteriaceae (**Fig 4A**). HOMINID can also detect SNPs where multiple taxa are correlated with the same SNP (e.g., SNP rs6032 in **Fig. 4B**), as well as more complex patterns of association; for example, for SNP rs230898 in the gene *TEKT3* (**Fig 4C**) genetic variation is positively correlated with one taxon (Clostridia) and negatively with others (Rhodocyclales and Aerococcaceae).

**Figure 4.**
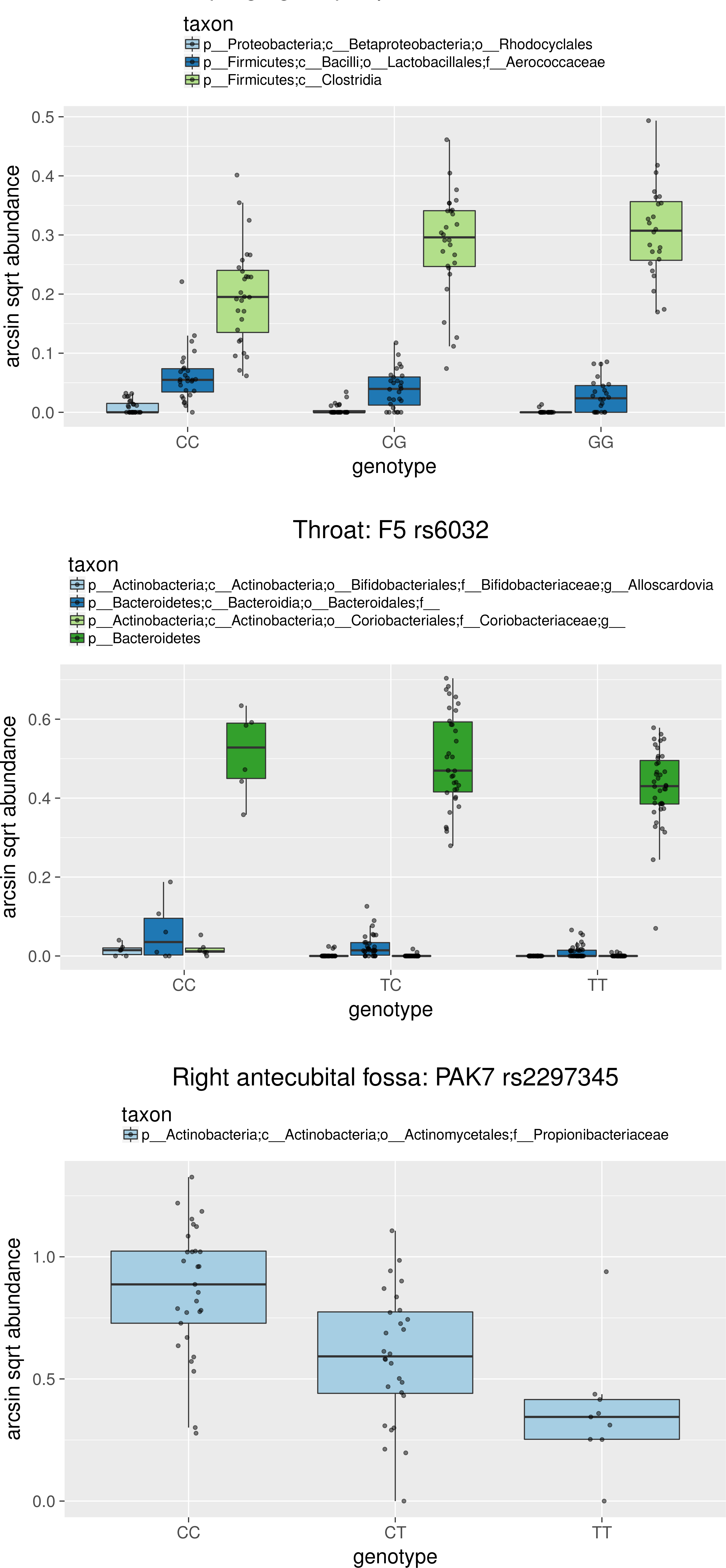
Examples of SNPs where correlations were found between host genetic variation and the microbiome. Three SNPs are shown: rs2297345 (correlated with abundance of microbial taxa in the right antecubital fossa), rs6032 correlated with abundance of microbial taxa in the throat), and rs230898 (correlated with abundance of microbial taxa in the supragingival plaque). The x-axis shows the host SNP genotypes, and the y-axis shows the arcsin sqrt transformed taxon abundances. The different correlated taxa for each SNP are shown in different colors.

Although HOMINID performs strongly on the data used in this paper, there are several potential limitations to our method. First, since it is especially designed to identify SNPs where a number of taxa are associated, it might not be optimal for cases where there is a dramatic shift in the microbiome that includes many dozens of taxa. Moreover, since the SNP is used as the response in the HOMINID model, it is difficult to identify epistatic effects, whereby genetic variation in two or more loci interact to affect microbiome composition. Although HOMINID could still be used to detect these interactions, by including all genotype combinations as response variables; however, multiple hypothesis testing could be an issue, especially for microbiome association studies, where samples sizes are currently small relative to GWAS of other complex traits. Nevertheless, HOMINID might be useful for detection of interaction of between candidate loci.

Lastly, we developed a web-based tool for the visualization of host-microbiome interaction network identified in HOMINID, available at http://z.umn.edu/genemicrobe. The website, designed using D3.js with a dedicated MySQL database serving as the back-end, displays a dynamic visualization of host gene-microbiome taxa interaction networks, and allows the user to add and remove nodes (host gene and microbial taxa), adjust the display size and node locations, filter by body sites, and generate figures. Currently, the website includes toy data representing all SNP-microbe associations with a nominal p-value <= 0.1 in the Human Microbiome Project data described above. We believe that as studies using larger sample sizes materialize (for example, a recent study included 1,514 subjects (Bonder et al. 2016)), we expect this tool to be useful for visualization of much larger number of associations.

## Conclusions

We present HOMINID, a framework designed for identifying associations between host genetic variation and microbiome composition. We analyze synthetic data to show HOMINID’s overall strong performance, identify specific factors that may affect it, highlight HOMINID’s unique features, and show HOMINID’s utility with a real dataset. We expect that HOMINID would be useful for studies attempting to characterize the genetic basis of host-microbiome interactions.

